# Genetic variation in male mate choice for large females in *Drosophila melanogaster*

**DOI:** 10.1101/2024.10.05.616829

**Authors:** Grace S. Freed, Isabella G. Martinez, Avigayil Lev, Ana-Maria Anthony Cuadrado, Alison Pischedda

**Affiliations:** Department of Biology, Barnard College, Columbia University, New York, NY

**Keywords:** male mate choice, body size, sexual selection, heritability, hemiclonal analysis

## Abstract

Males in many species show courtship and mating preferences for certain females over others when given the choice. One of the most common targets of male mate choice in insects is female body size, with males preferring to court and mate with larger, higher-fecundity females and investing more resources in matings with those females. Although this preference is well-documented at the species level, less is known about how this preference varies within species and whether there is standing genetic variation for male mate choice within populations. We used hemiclonal analysis in the fruit fly, *Drosophila melanogaster*, to test for heritable genetic variation in pre- and post-copulatory components of male mate choice for large females. We found additive genetic variation for both forms of male choice: males from different hemiclone lines varied in the strength of their courtship preferences for large females and the degree to which they extended matings with large females. Although males from hemiclone lines with stronger courtship preferences for large females were more likely to mate with those females, there was no genetic correlation between pre- and post-copulatory components of male mate choice, suggesting that they are under independent genetic control. Genetic variation in male mate choice may be widespread, potentially impacting the fitness of both sexes and the adaptive evolution of populations.

## INTRODUCTION

Females are typically thought of as the choosy sex, but male mate choice is a common phenomenon in many species (Bonduriansky 2001, Edward and Chapman 2011). Pre-copulatory male mate choice occurs when males preferentially court or mate with certain females over others (Edward and Chapman 2011), whereas post-copulatory cryptic male mate choice can manifest as differences in mating duration (Anastasio et al. 2023), ejaculate allocation (Wigby et al. 2009), and/or mate guarding (Ancona et al. 2010). Male mate choice is frequently directed toward higher fecundity females that can provide direct fitness benefits to males by producing more offspring (Bonduriansky 2001, Edward and Chapman 2011).

Male mate choice is often studied at the species level, such that any documented male preferences represent species-wide averages. In comparison, the potential for variation in male mate choice within species has received considerably less attention. Of the studies that have tested for intraspecific variation, most have compared male mate choice between populations with different life histories (Edward and Chapman 2013a) or between males that differ in phenotypic quality (Pollo et al. 2022), age (Edward and Chapman 2013b), or mating history (Dukas and Dukas 2012, Balaban-Feld and Valone 2018, Sinclair et al. 2021). For instance, males will sometimes modify their preferences based on their past mating experiences, increasing courtship toward types of females they have been successful with and decreasing courtship towards types of females they have been rejected by (Dukas and Dukas 2012, Balaban-Feld and Valone 2018). Additionally, smaller or lower quality males in a population can exhibit weaker preferences for high quality females (Pollo et al. 2022) and sometimes even prefer lower quality females (Baldauf et al. 2013, Pollo et al. 2019). It is unclear, however, whether any of this phenotypic variation in male mate choice has a genetic basis. If populations harbor standing genetic variation in male mate choice, as is often found for female mate choice (e.g., Jennions and Petrie 1997, Gray and Cade 1999, Rodríguez et al. 2013, Kelly 2018), this could impact sexual selection, the evolution of mutual mate choice (Servedio and Lande 2006, Courtiol et al. 2016), and the heritability of fitness across generations (Pischedda and Chippindale 2006, Bonduriansky 2009, Long et al. 2009).

One of the most common targets of male mate choice is female body size (Bonduriansky 2001, Edward and Chapman 2011), which is positively correlated with fecundity in most ectothermic species (Honěk 1993, Roff 2002). Pre- and post-copulatory male preferences for large females are especially well-documented in insects (Bonduriansky 2001), where males direct more courtship and mating efforts toward large females (Xu and Wang 2009), mate for longer with large females (Parker et al. 1999, Engqvist and Sauer 2003), and/or transfer more sperm and larger ejaculates when mating with large females (Gage and Barnard 1996, Gage 1998). In the fruit fly, *Drosophila melanogaster*, males prefer to court and mate with larger females (Byrne and Rice 2006, Sinclair et al. 2021), mate for longer with large females compared to small females (Lefranc and Bundgaard 2000, Lüpold et al. 2011, Anastasio et al. 2023), and may transfer more sperm and ejaculate when mating with larger females (Lüpold et al. 2011, Wigby et al. 2016). Despite these species-wide patterns, there is variation in the strength and direction of male mate choice between populations of *D. melanogaster* (Edward and Chapman 2013a), and between individuals from the same population (Byrne and Rice 2006, Sinclair et al. 2021, Lev and Pischedda 2023). The biological basis for this variation within *D. melanogaster* remains unclear, though it is likely not based on a male’s previous mating experience (Sinclair et al. 2021) or body size (Lev and Pischedda 2023).

We sought to investigate whether this variation in male mate choice for large females has a genetic basis in *D. melanogaster*. This hypothesis has not yet been tested in this species (or in any insect species, to the best of our knowledge), but there is genetic variation for both pre-and post-copulatory female mate choice in *D. melanogaster* (Giardina et al. 2011, Ratterman et al. 2014, Filice and Long 2017) and males frequently exhibit genetic variation in reproductive traits that may be associated with male mate choice, such as courtship intensity (Ruedi and Hughes 2008), mating speed (Veuille and Mazeau 1988, Hoffmann 1999), mating duration (Gromko 1987, Veuille and Mazeau 1988, Gaertner et al. 2015), and traits associated with ejaculate transfer (Chow et al. 2010, Pischedda et al. 2011). In addition, there is genetic variation in male mate choice based on female cuticular hydrocarbon pheromones in *D. melanogaster* (Pischedda et al. 2014), although genetic variation was not found when males were allowed to choose amongst female genotypes of unknown quality (Ratterman et al. 2014). If the direct benefits to males of mating with larger females are consistently strong in a population, any variation in male mate choice may have been eroded, such that all males show a strong preference for large females. In contrast, if the benefits to males are inconsistent, we might expect to see standing heritable variation reflecting weaker selection and/or alternative male approaches to mate choice.

In this study, we tested for heritable variation in male mate choice in *D. melanogaster* using hemiclonal analysis, a technique in which haploid genotypes are sampled from a population, replicated, and expressed in multiple males carrying random genetic backgrounds from the same population (Rice et al. 2005). We set up male mate choice trials where individual males from 29 hemiclone lines were paired with a large female and a small female. We used the data from these trials to measure several traits associated with male mate choice, including male courtship and mating preferences (components of pre-copulatory male mate choice) and mating duration (a component of post-copulatory cryptic male mate choice). This experimental design allowed us to test for additive variation in male mate choice and for genetic correlations between different components of male mate choice (Rice et al. 2005, Abbott and Morrow 2011).

Understanding the genetic basis of male mate choice and the relationship between its components will provide insight into its capacity to act as a target for adaptive evolution.

## METHODS

### *Drosophila* population maintenance

This experiment used *D. melanogaster* flies from the LH_M_ population, a detailed description of which can be found elsewhere (Rice et al. 2005). LH_M_ is a large, outbred, wild-type population that had been held under consistent laboratory conditions for over 700 generations at the time of our experiments. This population is maintained on a two-week, discrete lifecycle in 25 mm diameter vials containing 5-10 mL of food medium made up of cornmeal, molasses and yeast. In each generation, on day 12 post-egg deposition, flies from 56 vials are mixed, lightly anesthetized with CO_2_, and distributed among 56 new vials containing food medium supplemented with 6.4 mg of live yeast with 16 males and 16 females per vial (1792 breeding individuals). After 2.5 days, these adult flies are transferred into fresh unyeasted vials for an 18-hr period and then discarded, at which point excess eggs are manually culled to a density of 150–200 eggs per vial to begin the next generation. All flies, including those used for experiments, were incubated under a 12h:12 h light-dark cycle at 25 °C and 50–70% humidity.

### Creating hemiclone lines and collecting experimental hemiclone males

A full description of the cytogenetic cloning protocol used to create hemiclone lines in *D. melanogaster* can be found in Rice at al. (2005). Briefly, we use this technique to isolate haploid genomes containing chromosomes I (X), II, and III (accounting for >99% of a complete haplotype) from the LH_M_ population. We then replicated and maintained these haploid genomes using specialized “clone generator” females that enforce cosegregation and intact transmission of the three chromosomes from father to son (facilitated by the lack of molecular recombination in Dipteran males). To express these haploid genotypes in a wild-type background, these males were mated to wild-type LH_M_ females with a compound X chromosome (C(1)DX, *y*, *f*), thus creating wild-type hemiclone lines in which all males share the focal haplotype coupled with a random set of chromosomes from the LH_M_ population. We can use hemiclonal analysis, in which multiple individuals are phenotyped from several different hemiclone lines sampled from a population, to test for standing additive genetic variation in a trait of interest (Rice et al. 2005, Abbott and Morrow 2011). Statistical models that include hemiclone line as a random effect can then be used to estimate heritability (h^2^) for a trait as two times the additive genetic variation (i.e., the phenotypic variance attributed to hemiclone line) divided by the total phenotypic variance. The additive variation is multiplied by two here because individuals within a hemiclone line share half their genetic variation in common (Rice et al. 2005, Abbott and Morrow 2011). The hemiclone genome is always inherited intact without recombination, so it could contain epistatic variation between alleles within the haplotype. These effects are likely to be small, but the heritability estimates obtained from hemiclonal analysis should be considered an upper bound (Rice et al. 2005, Abbott and Morrow 2011).

This experiment tested for genetic variation in male mate choice using males from 29 hemiclone lines created from the LH_M_ base population. Virgin wild-type hemiclone males were collected using light CO_2_ anesthesia within 6 h of eclosion 9-10 days after egg deposition. The males were held in vials containing a small amount of food medium at a density of 5 males per vial for 3-4 days before the male mate choice experiments began. We set up male mate choice trials across 12 experimental blocks, with 10 randomly selected hemiclone lines tested per block, such that each hemiclone line was measured across 4-5 experimental blocks.

### Creating large and small females

We created the large and small females used in the male mate choice trials by manipulating larval density using the method established by Byrne and Rice (2006). We placed 16 male-female pairs of adult LH_M_ flies into egg collection cages (3.75 cm diameter by 5.8 cm high; Genesee Scientific) fitted with a Petri dish containing 5 mL of food medium. After ∼18 h, the adult flies were discarded, and eggs were randomly sampled from the Petri dishes and placed into unyeasted vials of fresh food. Large females were created by transferring 50 eggs into vials containing 10 mL of food medium, while small females were created by placing 100 eggs into vials containing 1 mL of food. Because development time is slower in the 1 mL vials, the eggs for the small females were collected 1 day in advance of the eggs for the large females to ensure the females could be collected at the same post-eclosion age. Vials were incubated under standard conditions for 9 or 10 days (for large and small flies, respectively), at which time females were collected from both treatments using light CO_2_ anesthesia within 6 h of eclosion to ensure virginity. Females were incubated in unyeasted vials containing a small amount of food medium at a density of 10 females per vial for 3-4 days before experiments. Female virginity was confirmed before experiments by examining holding vials and discarding any containing larvae.

The large and small females produced using this approach have non-overlapping size distributions and are visually distinguishable, but have body sizes that fall within the naturally occurring range of female sizes in the LH_M_ population (Anastasio et al. 2023, Stewart et al. 2024). LH_M_ males readily court and mate with large and small females created using this approach and show comparable mate choice preferences for large females when female size is naturally sampled or experimentally induced (Long et al. 2009, Edward and Chapman 2013a, Sinclair et al. 2021, Anastasio et al. 2023). We can therefore use larval density manipulation as a high-throughput method for generating standardized variation in female size for male mate choice experiments.

### Male mate choice trials

Mate choice trials were conducted when all experimental flies were 3-4 days post-eclosion. The evening before the trials, one large and one small virgin female were transferred via mouth aspirator into vials containing a small amount of food medium and returned to the incubator overnight. Once the incubator lights came on the next morning, one hemiclone male (from one of the 10 randomly selected lines for that experimental block) was aspirated into each vial containing one large and one small female. Vials containing males from the 10 hemiclone lines were then equally distributed amongst observers (5-7 per block), with each observer watching 10 vials at a time in two back-to-back set ups, such that each observer watched a total of 20 vials per experimental block. A foam plug was pushed down into each vial until the flies had an interaction space with a height of ∼1 cm. During the male mate choice trials, each observer watched their 10 vials continuously for a period of 30 minutes and collected minute-by-minute courtship data. Courtship was noted for each minute in which a male performed a courtship song (i.e., extended a wing and vibrated it when near a female) or an attempted copulation, and the body size of the female that received the courtship (large or small) was also noted. If a male courted both the large and the small female within the same minute, both courtship events were reported. Courtship was recorded up to and including the minute at which the male began mating, or until 30 minutes had passed. Matings that began within this 30-minute period were watched to completion. Mating start and end times, as well as the size of the female mated (large or small) were reported for the male’s first mating only. All observations were completed within the first 2 hours of the light cycle. In total, we measured male mate choice for 40-49 males from each of 29 hemiclone lines.

### Data processing and analysis

We obtained several metrics of male mate choice from our experiments. To measure male pre-copulatory courtship preferences, we calculated a preference index (PI) for each male using the number of minutes the male spent courting each female (Sinclair et al. 2021, Lev and Pischedda 2023). The preference index (PI) is calculated as:

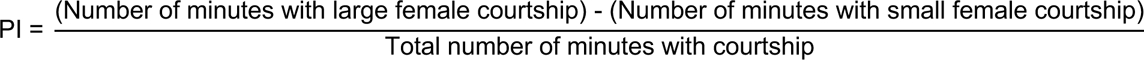

The PI can range from −1 to 1, with a negative PI indicating a preference for the small female, a positive PI indicating a preference for the large female, and a PI of 0 indicating no preference. The absolute value of the preference index indicates the strength of the preference, with PIs closer to 1 or −1 indicating a stronger preference than a PI closer to 0. We presented our results as PIs to be consistent with past male mate choice studies in LH_M_ (Sinclair et al. 2021, Lev and Pischedda 2023) to allow us to compare our findings to the population-wide averages reported in those studies. However, because of the negative values associated with PI data, they cannot be easily analyzed using the mixed model approaches necessary to estimate heritability. Therefore, to statistically analyze these courtship data, we converted PI to a binomial scale (i.e., with large female courtship as a success and small female courtship as a failure). We calculated the total proportion of courtship directed toward the large female in the following way: each courtship minute was considered two units (one for each female), each minute in which the male only courted the large female was assigned a score of 2, each minute in which the male courted both the large and the small female was assigned a score of 1, and each minute in which the male only courted the small female was assigned a score of 0. The proportion courtship directed to the large female obtained on this binomial scale (i.e. total large female score / total courtship units) correlates perfectly with our estimates of PI (r =1, proportion large female courtship = 0.5 + 0.5*PI) but enables us to use a binomial generalized linear mixed model (GLMM) to test for differences in male pre-copulatory preferences. We used *glmer* in the lme4 package (version 1.1.35.5; Bates et al. 2015) in R 4.4.1 (R Core Team 2024) to perform a binomial GLMM with hemiclone line, observer, and experimental block as random effects. We obtained the variance component estimates for each random effect and calculated 95% confidence intervals (CIs) on these variance components by profiling the likelihood surface using lme4. We then ran a parametric bootstrap test to obtain a p-value for the hemiclone line variance component using pbkrtest (version 0.5.3; Halekoh and Højsgaard 2014). To estimate h^2^ for preference index, we obtained the percent variance in PI attributed to hemiclone line using *r.squaredGLMM* in the MuMIm package (version 1.48.4; Bartoń 2024). We used the “theoretical” (distribution-specific) variances for all binomial models (Nakagawa et al. 2017).

We used courtship data (i.e., preference index) as our main measure of pre-copulatory male mate choice because metrics that involve mating are likely affected by female mate choice as well. However, matings are at least partly determined by male mate choice, so we also calculated the total proportion of matings with the large female (vs. the small female) for each hemiclone line. We tested for genetic variation in this trait using a binomial GLMM in *lme4* (with large female matings = 1 and small female matings = 0) with random effects hemiclone line and observer (block was not included in this model because it caused singularity issues). We also used a binomial GLMM to test for genetic variation in overall male mating success (i.e., the proportion of matings that occurred during our trials, with a successful mating = 1 and no mating = 0). For each model, we obtained the variance component estimates, the 95% confidence intervals and p-values for these estimates, and calculated h^2^, as described above.

We used mating duration as our measure of post-copulatory cryptic male mate choice, as this is primarily under male control in *D. melanogaster* (Jagadeeshan and Singh 2006, Friberg 2006). Although *D. melanogaster* males typically mate for longer with large females versus small females (Lefranc and Bundgaard 2000, Anastasio et al. 2023), it is unclear whether there is genetic variation in the degree to which males extend matings with larger females. To test for variation in this trait, we used *lmer* in lme4 to conduct a linear mixed model with the size of the female mated (large or small) as a fixed effect and the following random effects: hemiclone line, the hemiclone line-by-size of female mated interaction, observer, and experimental block. A significant interaction between hemiclone line and the size of the female mated would be consistent with genetic variation in post-copulatory male choice, as it indicates that hemiclone males varied in the degree to which they extended matings with large females. We used *ANOVA* in car (version 3.1.2; Fox and Weisberg 2019) to analyze the fixed effect female size and we used lme4 to obtain 95% confidence intervals on the variance component estimates for each random effect and obtained p-values for these estimates using pbkrtest. We the used MuMIm to estimate h^2^ for overall mating duration (using the variance attributed to hemiclone line) and post-copulatory male mate choice (using the variance attributed to the hemiclone-by-size of female mated interaction).

Finally, we tested for genetic correlations between components of male mate choice using JMP 18. For each hemiclone line, we obtained the following data: mean preference index, proportion of matings with large females, mating success (proportion of successful matings), and the relative mating duration extension with large females (i.e., mean mating duration with large females / mean mating duration with small females). We used Spearman’s rank correlation tests to check for genetic correlations between different components of male mate choice.

## RESULTS

### Genetic variation in pre- and post-copulatory male mate choice

There was significant variation in preference index between hemiclone lines (estimated h^2^=0.20, p=0.0012; Table 1a, Table 2), revealing standing additive genetic variation in the strength of male courtship preferences for large females. Males from our sample of 29 haplotypes ranged from having no apparent courtship preference between large and small females to a strong preference for large females (Figure 1). In contrast, we did not find significant variation in the proportion of matings that occurred with the large female among hemiclone lines (Table 1b, Table 2). Although not the focus of our study, we found substantial genetic variation in overall male mating success (estimated h^2^=0.42, p=0.0063; Table 1c, Table 2).

**Figure 1.**
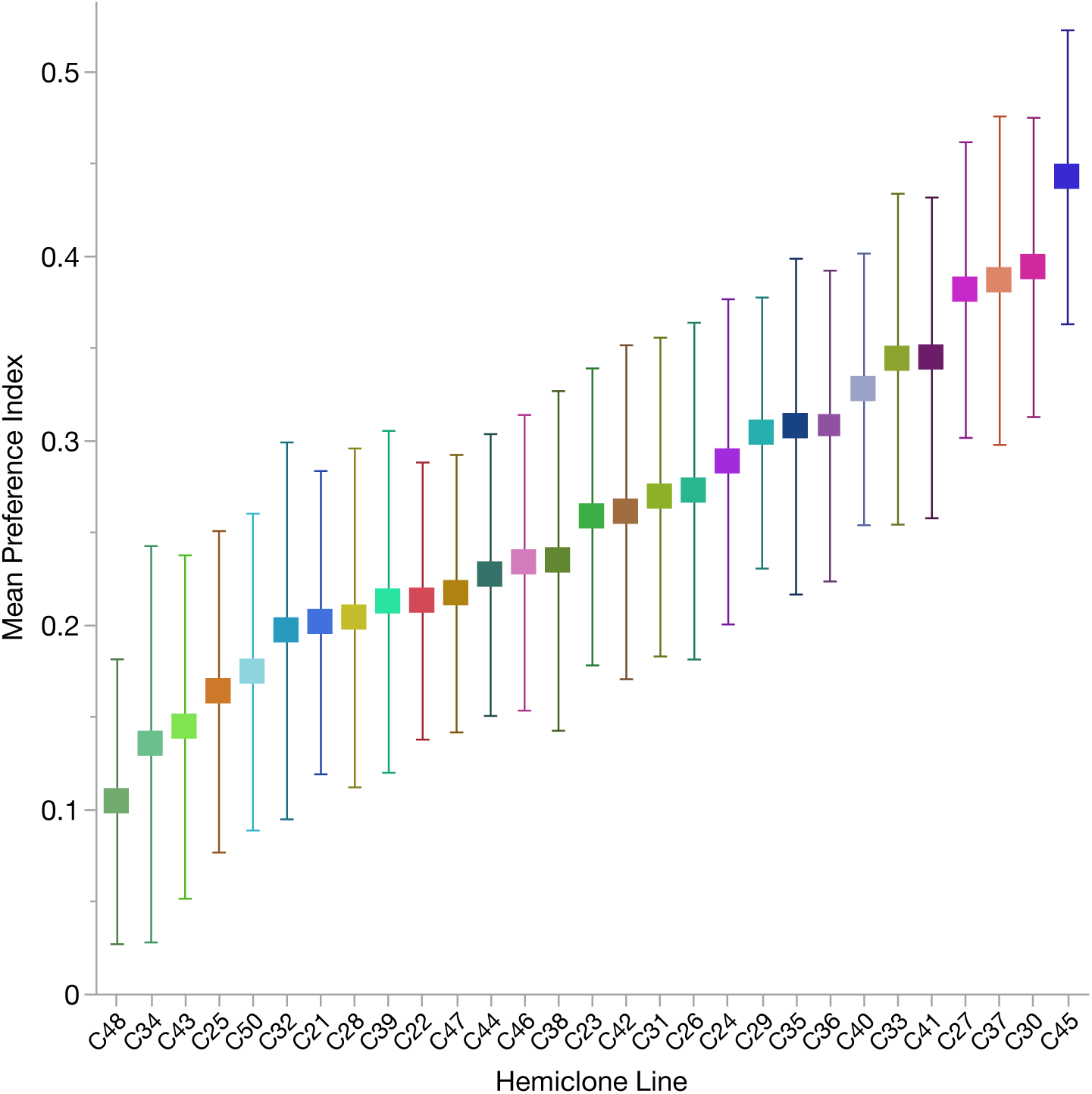
Mean preference index (± 1 standard error) for males from the 29 hemiclone lines surveyed in our study (N=40-49 males per hemiclone line), demonstrating standing additive variation in pre-copulatory male mate choice (h^2^=0.20).

**Table 1.**
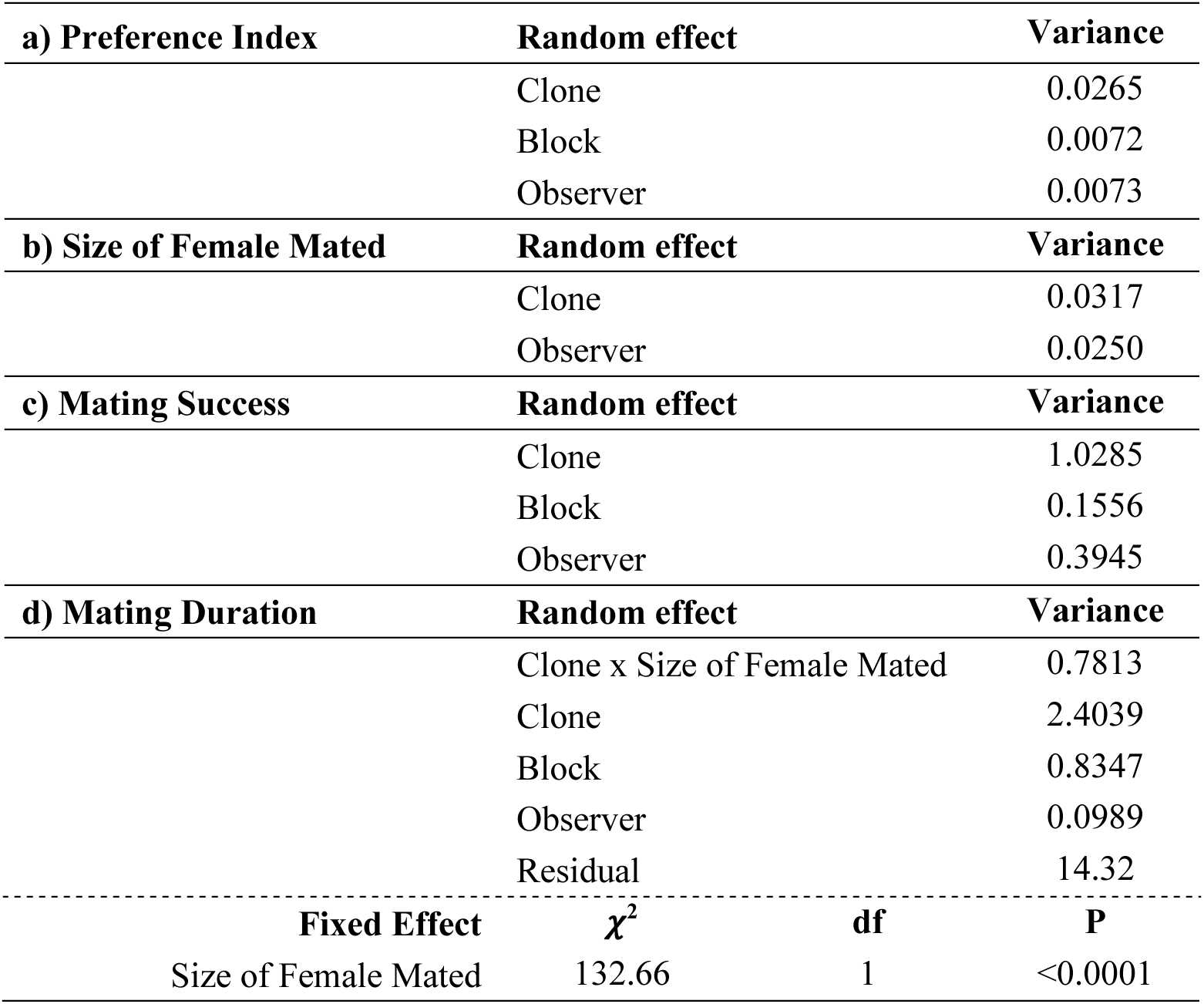
Results of the mixed models testing for variation in components of male mate choice. Shown are the results for: a) preference index, b) the size of the female mated, c) mating success, and d) mating duration.

**Table 2.**
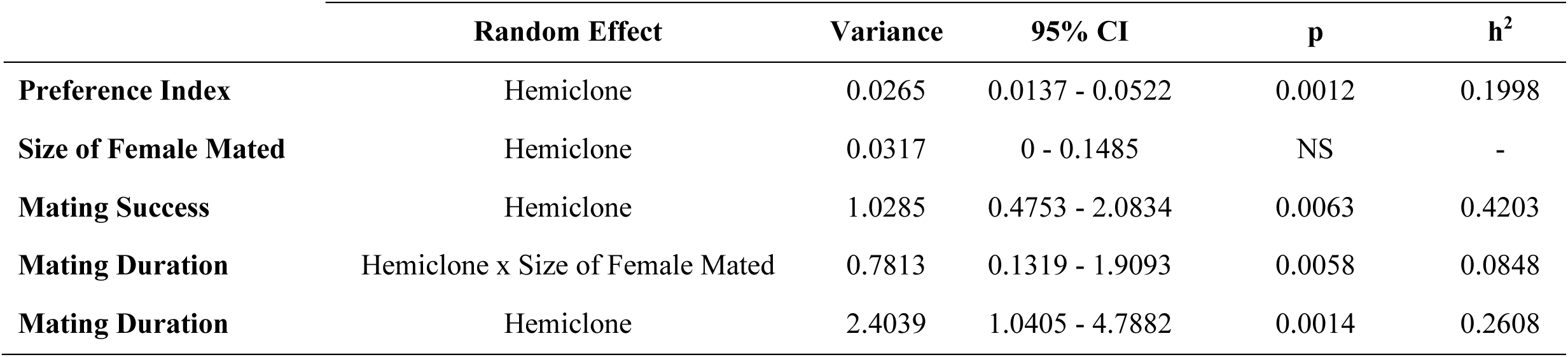
Hemiclone variance components for measures of male mate choice, including preference index, size of female mated, mating success, and mating duration. Shown for each phenotype is the hemiclone variance estimate (and the hemiclone x size of female mated variance estimate for mating duration), the 95% confidence intervals (CIs) and p-values associated with this estimate, and the estimated heritability (h^2^) for this phenotype. NS indicates that the p-value was not significant.

Once mating was initiated, hemiclone males mated for longer with large females than with small females, on average (p<0.0001, Table 1d), and there was standing additive variation in overall male mating duration (estimated h^2^=0.26, p=0.0014, Table 1d, Table 2). We also found additive variation attributed to the interaction between hemiclone line and the size of the female mated (estimated h^2^ = 0.08, p=0.0058, Table 1d, Table 2), indicating that hemiclone males varied in the degree to which they extended matings with large females (Figure 2).

**Figure 2.**
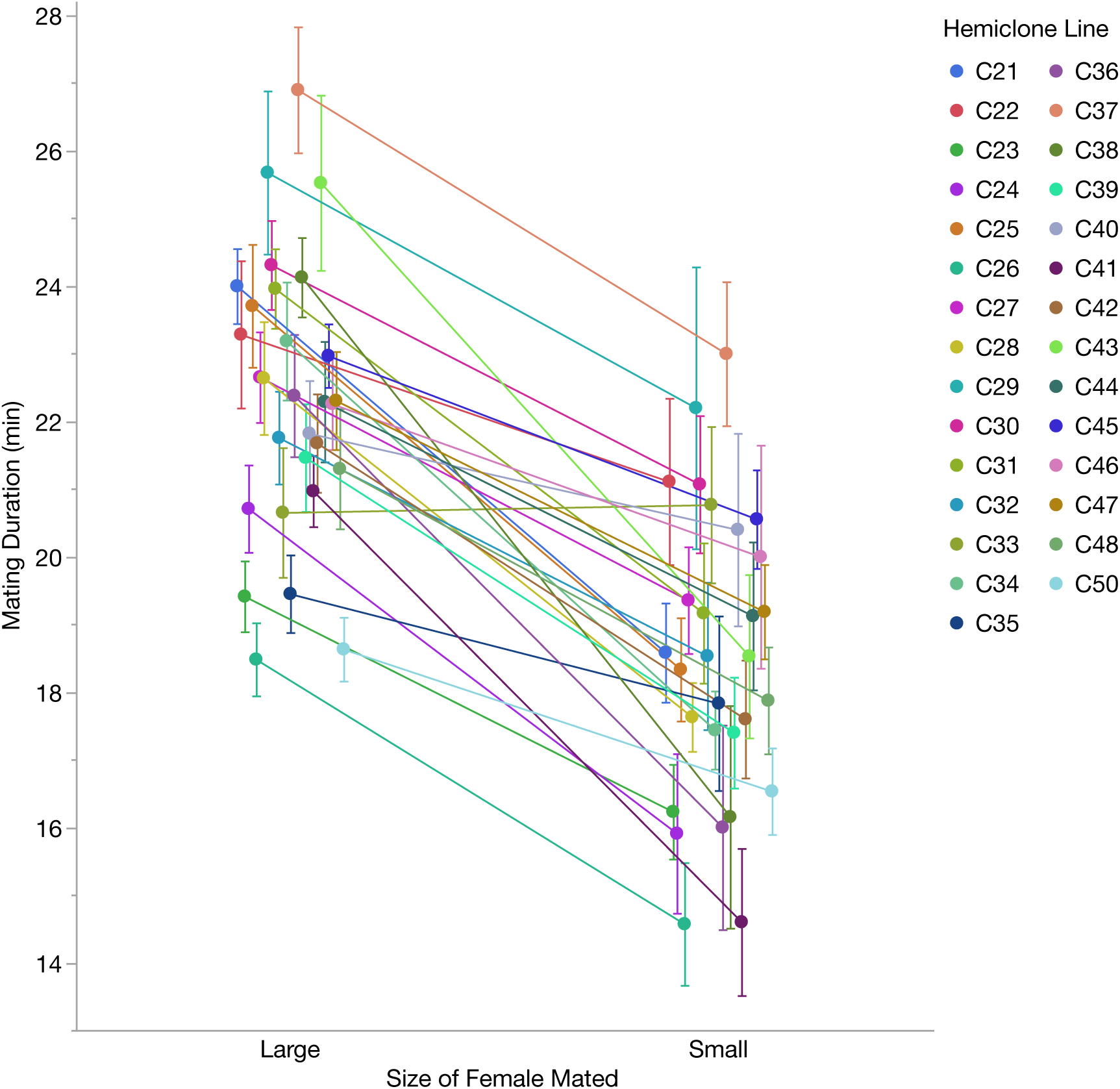
Mean mating durations (± 1 standard error) for males from 29 hemiclone lines when mating with either large females (N=16-36 males per hemiclone line) or small females (N = 5-21 males per hemiclone line). We found standing additive variation associated with overall mating duration (h^2^ = 0.26) and for the interaction between hemiclone line and the size of the female mated (i.e., our measure of post-copulatory male mate choice, h^2^ = 0.08).

### Genetic correlations between components of male mate choice

We found a significant genetic correlation between mean preference index and the proportion of matings with large females for our 29 hemiclone lines (r_s_= 0.59, p=0.0007, Figure 3a), indicating that haplotypes with stronger courtship preferences for large females were more likely to mate with those females. In contrast, we found no genetic correlation between preference index (our measure of pre-copulatory male mate choice) and the relative mating duration extension with large females (our measure of post-copulatory cryptic male mate choice), suggesting that pre- and post-copulatory components of male mate choice are likely independent at the genetic level (r_s_= −0.30, p=0.11; Figure 3b). Last, we tested for genetic correlations between mating success and male mate choice to determine if successful males were more likely to be choosy (Pollo et al. 2022), but we found no correlation between mating success and preference index (r_s_=0.20, p=0.29) or the relative mating duration extension with large females (r_s_= −0.26, p=0.18).

**Figure 3.**
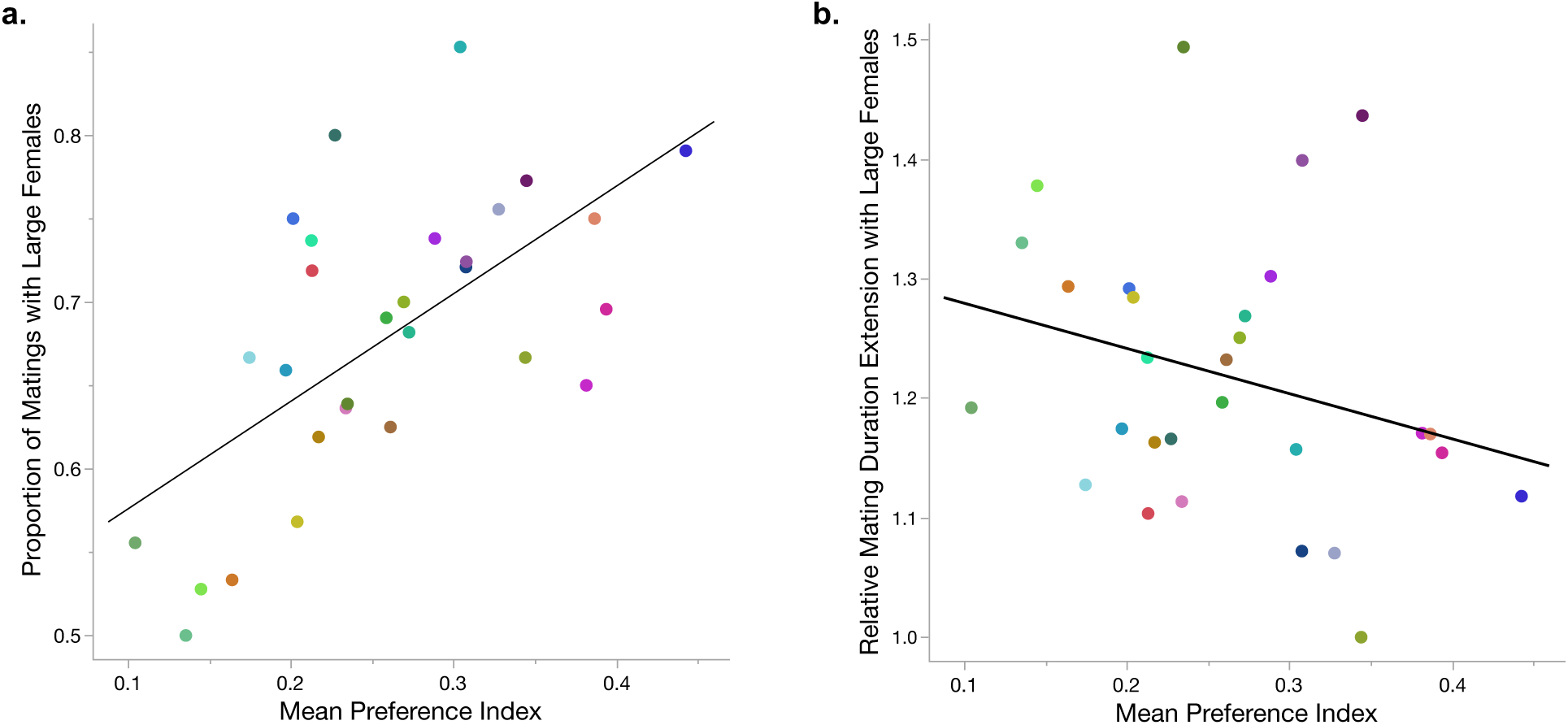
Correlations between components of male mate choice. a) Genetic correlation between mean preference index (our measure of pre-copulatory male mate choice) and the proportion of matings that occurred with the large female (vs. the small female) for males from 29 hemiclone lines (r_s_= 0.59, p=0.0007). b) No genetic correlation between mean preference index and the relative mating duration extension with large females (our measure of post-copulatory male mate choice, calculated as mean mating duration with large females / mean mating duration with small females) for males from 29 hemiclone lines (r_s_= −0.30, p=0.11). Each point for both a and b represents a specific hemiclone line; point colors are consistent with those used in Figures 1 and 2.

## DISCUSSION

Genetic variation in mate choice can influence the outcome of sexual selection, the evolution of reproductive traits and behaviors, and the adaptive evolution of a population (Jennions and Petrie 1997, Millan et al. 2020). Although heritable variation in female mate choice has been documented in many species (Jennions and Petrie 1997, Gray and Cade 1999, Giardina et al. 2011, Rodríguez et al. 2013, Ratterman et al. 2014, Filice and Long 2017, Kelly 2018), the potential for similar variation in male mate choice has received much less study. Using hemiclonal analysis in *D. melanogaster*, we found standing additive genetic variation in both pre- and post-copulatory male mate choice for large females, a common target of male mate choice in insects (Bonduriansky 2001).

The presence of genetic variation in preference index, our measure of pre-copulatory male mate choice, suggests that male genotype may contribute to the individual variation in male mate choice observed within populations (Byrne and Rice 2006, Sinclair et al. 2021, Lev and Pischedda 2023) and lead to changes in male preferences between populations with different life histories (Edward and Chapman 2013a). Notably, males in our experiment ranged from having no apparent courtship preference between large and small females to a strong preference for large females, with no hemiclone line showing a preference for small females (as indicated by a negative mean preference index, Figure 1). This is not surprising, as *D. melanogaster* males surveyed from the LH_M_ population typically have mean preference indexes ranging from approximately 0.2 to 0.5 (Sinclair et al. 2021, Lev and Pischedda 2023). Since most males spend more time courting large females in this population, any males with preferences for small females should be rare (if they exist at all). While the mean preference index for these 29 hemiclone lines (PI = 0.26) was within the normal range for this population, we would likely need to survey many more haplotypes to detect any potential male preferences for small females.

In contrast, we did not find additive variation in the proportion of matings that occurred with the large female. A successful mating with a large female will be influenced by both male and female mate choice, so the lack of variation in this trait highlights the complicated nature of partitioning mate choice between the sexes. Measuring courtship preferences can be challenging in many species, but our results suggest that measuring male mate choice using mating-related traits only (such as which female a male mated with) may fail to detect existing variation in male preferences. Nevertheless, we found a positive correlation between preference index and the proportion of matings with large females (Figure 3a), indicating that males with stronger courtship preferences for large females were, in fact, more likely to mate with those females.

We used mating duration as a measure of male investment and cryptic male mate choice, as it is primarily under male control in *D. melanogaster* (Jagadeeshan and Singh 2006, Friberg 2006). Mating duration is not associated with the number of sperm transferred in this species (Lüpold et al. 2011, Macartney et al. 2021), but longer matings may be energetically costly to males (Parker et al. 1999), take up time that could otherwise be spent securing additional mates (Daly 1978), and may deplete the males’ ejaculatory resources (Dewsbury 1982, Friberg 2006, Hopkins et al. 2019, Anastasio et al. 2023). We found genetic variation in overall male mating duration, as reported previously (Gromko 1987, Gaertner et al. 2015), but we also detected standing additive variation in the degree to which males extended matings with large females, our measure of post-copulatory cryptic male mate choice. As with our preference index data, no hemiclone line had males that mated for longer with small females (Figure 2). This may similarly be a result of our limited genetic sample, as *D. melanogaster* males typically mate for longer with large females than with small females on average (Lefranc and Bundgaard 2000, Anastasio et al. 2023).

Although we found standing genetic variation in male mate choice both before and after mating, the strengths of pre-copulatory and post-copulatory male mate choice were not genetically correlated in our study (Figure 3b). These components of male mate choice therefore act in the same direction (i.e., both show a mean preference for larger females), but appear to be under independent genetic control. It is possible that only surveying 29 haploid genomes limited our ability to detect an existing genetic correlation, but this genetic decoupling is consistent with other work showing that pre- and post-copulatory components of male fitness, including those influenced by female mate choice, are often not genetically correlated in *D. melanogaster* (Droge-Young et al. 2012, Travers et al. 2016).

Our study reveals additive variation in male mate choice for a common male preference in insects (Bonduriansky 2001), but it raises questions about why this variation persists. Choosier males were more likely to mate with larger, higher-fecundity females (Figure 3a), so selection should favor courtship preferences for large females in this population. If males similarly benefit from extending matings with larger females, for example, by delaying female remating (Gilchrist and Partridge 2000) and/or increasing male fertilization success (Friberg 2006), then higher post-copulatory male investment in large females should also be favored. When these fitness benefits to males are sufficiently strong, we might expect all males to have a strong preference for large females. However, if male mate choice creates stronger competition between males for larger females, less competitive males may have weaker preferences for females that they are less likely to be successful with (Edward and Chapman 2011). Despite male attractiveness contributing to variation in male mate choice in other species (Pollo et al. 2022), the lack of genetic correlations between mating success and either component of male mate choice found here suggests that this is not the case in this system.

It is possible that the direct fitness benefits associated with this male preference are not as strong as expected. We tested male mate choice in a non-competitive environment, so the correlation between pre-copulatory male mate choice and the likelihood of mating with large females may be weaker when other males are present. Additionally, in mating systems that experience sexual conflict, like *D. melanogaster*, male mate choice for large females can cause those females to receive a disproportionate amount of male harassment and/or harmful matings (Fowler and Partridge 1989, Partridge and Fowler 1990, Long et al. 2009). This bias in male harm is costly to large females and can minimize the fecundity differences associated with female size (Long et al. 2009). When males vary in the strength of their preference for large females, as shown here, they could also vary in the degree of harm they inflict on those females. If the choosiest males are most harmful to large females, this could lessen the direct fitness benefits received by those males and weaken selection on male mate choice. Even if large females do maintain higher fecundity, large females remate more rapidly than small females (Wigby et al. 2016, Anastasio et al. 2023), so there is no guarantee that a choosy male will receive all of the potential fitness benefits associated with female size. Measuring lifetime fitness for hemiclone lines that vary in their strength of male mate choice could help to improve our understanding of how selection shapes male preferences.

Male mate choice for large females in this laboratory-adapted population of *D. melanogaster* operates similarly to how it does in other insect species. Males direct more courtship and mating effort toward large females (Xu and Wang 2009, Sinclair et al. 2021), mate for longer with large females (Parker et al. 1999, Anastasio et al. 2023) and may transfer more sperm (Gage and Barnard 1996, Lüpold et al. 2011) and/or larger ejaculates (Gage 1998, Wigby et al. 2016) to larger females. Given these similarities, the additive variation in male mate choice we report here may be widespread across insects that share this male preference. We show that genetic variation in mating preferences is not unique to females, highlighting the similarities in mate choice evolution between the sexes. This variation has unexplored implications for the fitness benefits that males receive from being choosy, the consequences of this choosiness for females, and the potential for adaptive evolution at the population level.

## ACKNOWLEDGEMENTS

We thank Ashley Canizares, Ananya Lahiri, Olivia Anastasio, Chelsea Sinclair, and Abigail Gutierrez for help with the male mate choice experiments. We are grateful to Ian Dworkin and the members of the McMaster University Data Lunch seminar for help with our data analysis. This work was funded in part by a Barnard College Presidential Research Award to AP. GSF and IGM received additional support from Barnard’s Summer Research Institute, while AAC received support from a Con Edison Summer Grant.

## AUTHOR CONTRIBUTIONS

GSF: conceptualization (supporting), investigation (lead), visualization (supporting), writing – original draft preparation (equal)

IGM: conceptualization (supporting), investigation (lead), visualization (supporting), writing – original draft preparation (equal)

AL: investigation (supporting), writing – original draft preparation (equal), writing – review & editing (equal)

AAC: conceptualization (supporting), methodology (supporting), investigation (supporting), writing – original draft preparation (supporting)

AP: conceptualization (lead), methodology (lead), investigation (supporting), formal analysis (lead), visualization (lead), writing – original draft preparation (equal), writing – review & editing (equal), funding acquisition (lead)

